# Single-egg Comet Assay: a protocol to quantify DNA damage in aquatic dormant stages

**DOI:** 10.1101/2024.08.06.606806

**Authors:** Rejin Salimraj, Alessio Perotti, Marcin W. Wojewodzic, Dagmar Frisch

**Affiliations:** Pharmacovigilance, United BioSource Corporation, Geneva, Switzerland; School of Biosciences, The University of Birmingham, Birmingham, B15 2TT, United Kingdom; Department of Chemical Toxicology, Norwegian Institute of Public Health, Oslo, Norway; Department of Research, Cancer Registry of Norway, Norwegian Institute of Public Health, Oslo, Norway; Department of Evolutionary and Integrative Ecology, Leibniz Institute of Freshwater Ecology and Inland Fisheries (IGB), Berlin, Germany

**Keywords:** bioarchive, DNA degradation, sedimentary propagules, paleogenomics, resurrection ecology, *Daphnia*

## Abstract

1. The comet assay (CA) was originally developed as toxicity test and quantifies DNA integrity from the distribution of DNA across an electric field. Compromised DNA moves across electric fields faster than intact DNA strands, leaving a quantifiable signature that resembles a comet tail. The dimensions of this comet tail reflect relative DNA damage.
2. We optimized the CA protocol for individual dormant propagules (Single-egg Comet Assay or SE-CA) to inform downstream analyses such as DNA sequencing, of the DNA quality contained in natural genetic archives of past populations. As a model we used dormant eggs of the microcrustacean *Daphnia*.
3. We tested the SE-CA protocol on impact of processing and storage conditions for dormant eggs and used it to assess DNA damage related to aging of eggs retrieved from recently deposited to centuries-old lake sediment. The SE-CA successfully determined the degree of DNA damage in individual eggs frozen in liquid nitrogen, or at -80°C as well as damage caused by bleaching and historical egg age.
4. In conclusion, our protocol provides a cost-effective method of assessing DNA damage in sedimentary propagules such as dormant *Daphnia* eggs. More generally, the SE-CA can be applied to test DNA integrity in individual propagules prior to genome sequencing or to quantify environmental impacts on natural sedimentary biobanks.

## Introduction

The Comet Assay (CA) was originally developed as a toxicological test for human cells and can assess DNA damage in small numbers of cells (Ostling & Johanson, 1984; Collins *et al*., 2023). The CA is an OECD endorsed method for measuring genotoxicity of compounds (OECD, 2016) which, by inducing strand breaks at sites of damage such as alkali-labile sites and abasic sites, can accurately quantify both single and double strand DNA damage (Calini, Urani & Camatini, 2002). Development of the CA yielded many advantages, such as the lower cell count (less than 10,000 cells) required per sample compared with other techniques when assessing DNA damage (Collins *et al*., 1997; Dixon *et al*., 2002). In addition to human tissues (Jałoszyński *et al*., 1997), the CA is applied to study DNA degradation in invertebrate taxa (Gajski *et al*., 2019), e.g. after exposure to pollutants (Mitchelmore & Chipman, 1998), or in archived tissues and specimens (Raxworthy & Smith, 2021). The CA was also applied to the microcrustacean model *Daphnia*, to test the eco-genotoxicity of known pollutants such as hydrogen peroxide on DNA damage (Pellegri, Gorbi & Buschini, 2020) or to measure global methylation levels with the methylation sensitive CA (Kusari *et al*., 2017). However, to our knowledge the CA has never been applied to dormant eggs of aquatic taxa archived in lake sediments. These are of particular interest in ecological and evolutionary studies that use dormant stages contained in these natural archives for genetic analysis (Orsini *et al*., 2013; Ellegaard *et al*., 2020) or for their “resurrection” hatched from strata as old as centuries for evolutionary studies (Frisch *et al*., 2014; Yousey *et al*., 2018; Weider, Jeyasingh & Frisch, 2018). Well-preserved dormant *Daphnia* eggs from lake sediment up to 1000 years old have been studied genetically (Mergeay, Verschuren & De Meester, 2006; Brede *et al*., 2009; Frisch *et al*., 2016) and are also suitable for genomic applications (Lack, Weider & Jeyasingh, 2018; O’Grady *et al*., 2022).

We aimed to adapt and optimise a CA protocol for such dormant stages using eggs of *Daphnia* as a model. If successful, the low-cost CA could be applied to assess DNA quality in a number of dormant eggs from given sediment strata prior to pursuing time-intensive attempts to resurrect other isolates, or prior to whole genome sequencing for population genomic studies. In addition, its use as a test of the impact of pollution or other environmental impacts on natural biobanks would be desirable. Further, we evaluated the suitability of the here developed standardised SE-CA protocol by testing it on three potential sources of DNA degradation: (1) cryostorage (freezing at -80 °C, freezing in liquid nitrogen), (2) bleaching as a treatment for removal of exogenous DNA on egg surfaces prior to genomic applications, and (3) egg age across six centuries.

## Materials and Methods

### Daphnia egg collection and extraction

Days-old ephippia were isolated from a lab culture of *Daphnia magna* Straus, 1820, maintained at the University of Birmingham, UK. Historical Arctic *Daphnia pulicaria* Forbes, 1893 eggs with an estimated age range of the past ∼600 years (Dane *et al*., 2020) were isolated from sediment collected from lake SS4 in Kangerlussuaq, South-West Greenland. Eggs were removed from individual ephippia in phosphate-buffered saline (1X PBS, pH ∼7.4) and stored in 1X PBS at 5°C until further processing.

### Protocol for single-egg Comet Assay

Active lysis solution was prepared by adding 10% DMSO and 1% Triton-X 100 to the lysis stock solution and left to chill at 4°C. Individual fresh, days-old *Daphnia magna* eggs were homogenized in an Eppendorf tube with a pipette tip in a final concentration of 1% DMSO and 0.1mM Na_2_EDTA dissolved in 1X PBS.

Slides were pre-coated with normal melting point agarose (NMA) 2% (Promega, Southampton, UK). Each Eppendorf tube containing a homogenised egg was centrifuged at 3500 RPM for 90 seconds and the supernatant was carefully removed from the pellet produced. The pellet was resuspended with 20-25μl of previously prepared 0.5% low melting point agarose (LMA) preheated at 37°C and applied onto the precoated NMA slide which was then covered using a 22×22mm coverslip. The slides were left to set on a metal block at 4°C for 15 minutes and once hardened, the coverslips were removed carefully. After adding another layer of 20-25μl LMA, coverslips were replaced and the slides left to set at 4°C for another 15 minutes. After careful removal of the coverslips, slides were incubated in ice-cold lysis buffer (2.5M NaCl, 0.1M Na_2_EDTA, 10mM Tris) at 4°C. On the following day, electrophoresis was carried out in the dark. The slides stayed submerged in electrophoretic buffer (300mM NaOH, 1mM Na_2_EDTA) for 10 minutes to allow unwinding of the DNA coils before applying the current (0.78V/cm and 300mA) with constant voltage during electrophoresis. After electrophoresis, slides were neutralised with 0.4 M Tris-HCl buffer, adjusted to pH 7.5, fixed in absolute ethanol for ten minutes, stained with 1:10,000 dilution of SYBR-GOLD (Life Technologies ltd, Paisley, UK) and kept in a damp chamber to prevent drying.

### Quantification of DNA damage

Slides were examined under a fluorescent microscope (Zeiss, Axiovert) equipped with a 515 to 560 nm excitation filter and a barrier filter of 590 nm using a 40 X oil immersion lens. The level of DNA damage in each cell was determined using the Comet Assay IV software (Perceptive Instruments, Bury St Edmunds, UK) which detects relative fluorescent light intensity between intact (head) and fragmented DNA (tail). The light intensity of fluorescent fragmented tails is calculated by the Comet Assay IV software as a percentage of the total light intensity of both the head and the tail. Statistical analysis used tail intensity % as response variable, which has a linear relationship with DNA fragmentation (Azqueta & Collins, 2013).

### Single-egg Test

To test the replicability of the single-egg CA, four individual slides with one *D. magna* egg each were prepared and at least 50 nuclei were scored per egg. Data from this test was also used as control for the freeze assay and the bleaching assay.

### Freeze Assay

*Daphnia magna* eggs were stored in separate Eppendorf tubes in 16 μl of 1X PBS each. Two tubes were frozen for ten minutes either in a -80°C freezer or flash-frozen in liquid nitrogen (LN) and then left at room temperature to thaw followed immediately by CA for eggs from both tubes. For the control, four eggs were scored (from *Single-egg test*), two for LN and one egg for the -80°C treatment, scoring at least 50 nuclei per egg.

### Bleaching Assay

The DNA damage caused by bleaching was tested by comparing two types of eggs: days-old dormant *D. magna* eggs and sedimentary Arctic *D. pulicaria* eggs obtained from Braya Sø dating to *ca*. 1805 AD. Two eggs of each of *D*.*magna* (from *Single-egg test*) and *D. pulicaria* were used as control and two were treated with bleach solution. Eggs were exposed to a solution of 5% bleach solution made from industrial strength (12%) sodium hypochlorite for 30 seconds, then washed in PBS twice to remove any remaining bleach. Controls were untreated prior to CA.

### Age Assay

For this test, eggs were extracted from four sediment depths of Braya Sø, Southwest Greenland, representing (at the time of sampling) a range in age from ∼40 to ∼630 years old (*ca*. 1976 AD, *ca*. 1940 AD, *ca*. 1805 AD, *ca*. 1390 AD). For each time period, one egg was used for analysis of which at least 50 nuclei were scored.

### Statistical Analysis

To test for significant effects of the experimental conditions, we used tail intensity % as response variable in non-parametric Kruskal Wallis tests. This test was chosen because of the non-normal distribution of the response variable. Statistical significance between pairwise comparisons of treatments was computed with the Dunn test (R package dunn.test (Dinno, 2024)), and p-values adjusted with the Benjamini-Hochberg procedure for multiple testing. All analyses were computed on the R platform (R CoreTeam, 2017). Data was visualised with ggplot2 version 3.3.2 (Wickham, 2016).

Upon acceptance, the data and R code data will be made available in Zenodo.

## Results

To evaluate the application of the SE-CA, we tested the impact of four potential sources of DNA damage: freezing at -80°C and in liquid nitrogen, bleaching for removal of external DNA, and age of eggs from sediment archives (Fig. 1).

**Fig. 1.**
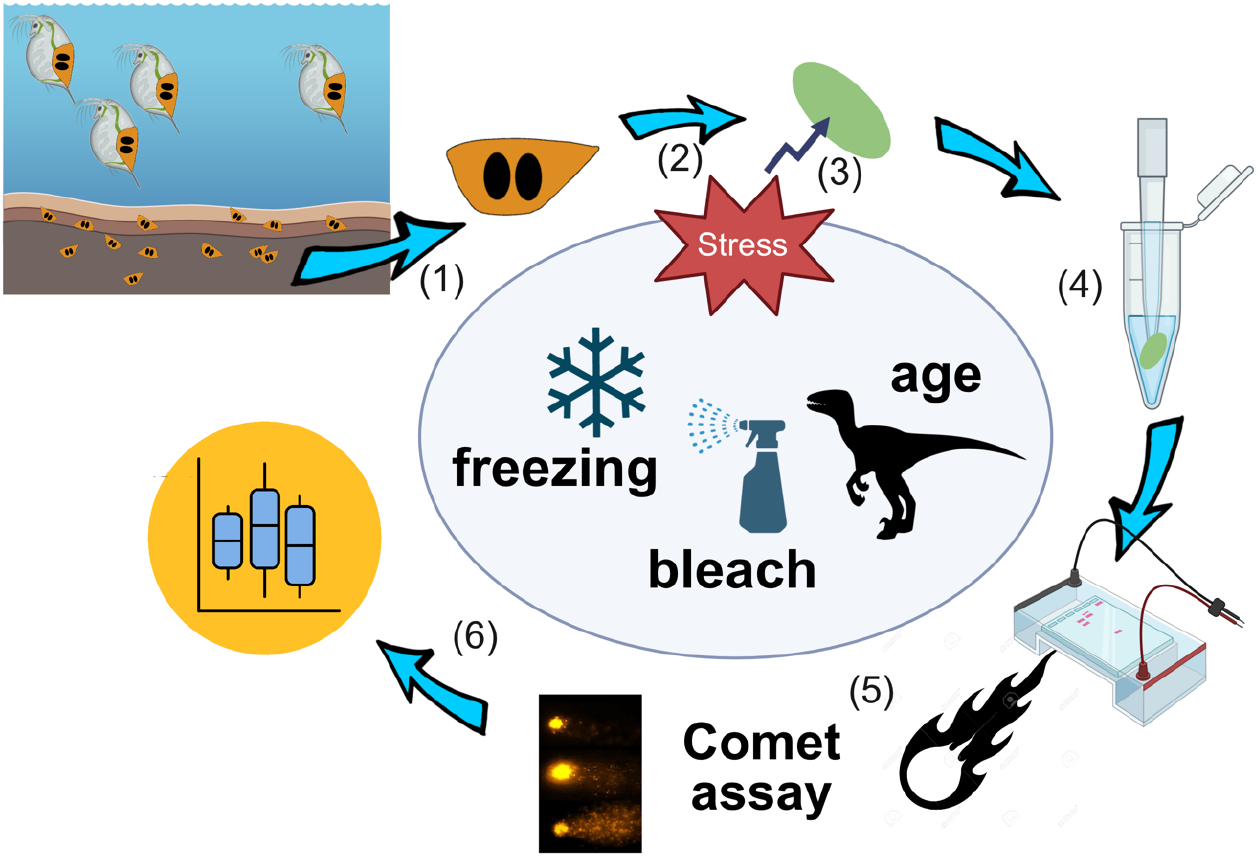
Workflow for the SE-CA on propagules from natural sedimentary bioarchives: (1) Isolation of sedimentary stages from sediment archive, (2) decapsulation of eggs (or other propagules), (3) artificial or natural stress impact (here: freezing, bleaching, aging), (4) homogenisation with pipette, (5) performance of comet assay and fluorescent microscopy according to protocol, (6) data analysis. For details see Methods. Figure created with BioRender.com.

### Single-egg test

We tested the reproducibility of the SE-CA protocol on individual *Daphnia magna* dormant eggs. DNA damage differed only slightly between four eggs individually processed on separate slides with median % tail intensity between 0.94 and 1.39% (Fig. 2a, Table S1). Although the Kruskal-Wallis test was significant (Table S2, p = 0.04), the only significant pairwise comparison was between slide 1 and 2 (p=0.047, Table S3)

**Fig. 2.**
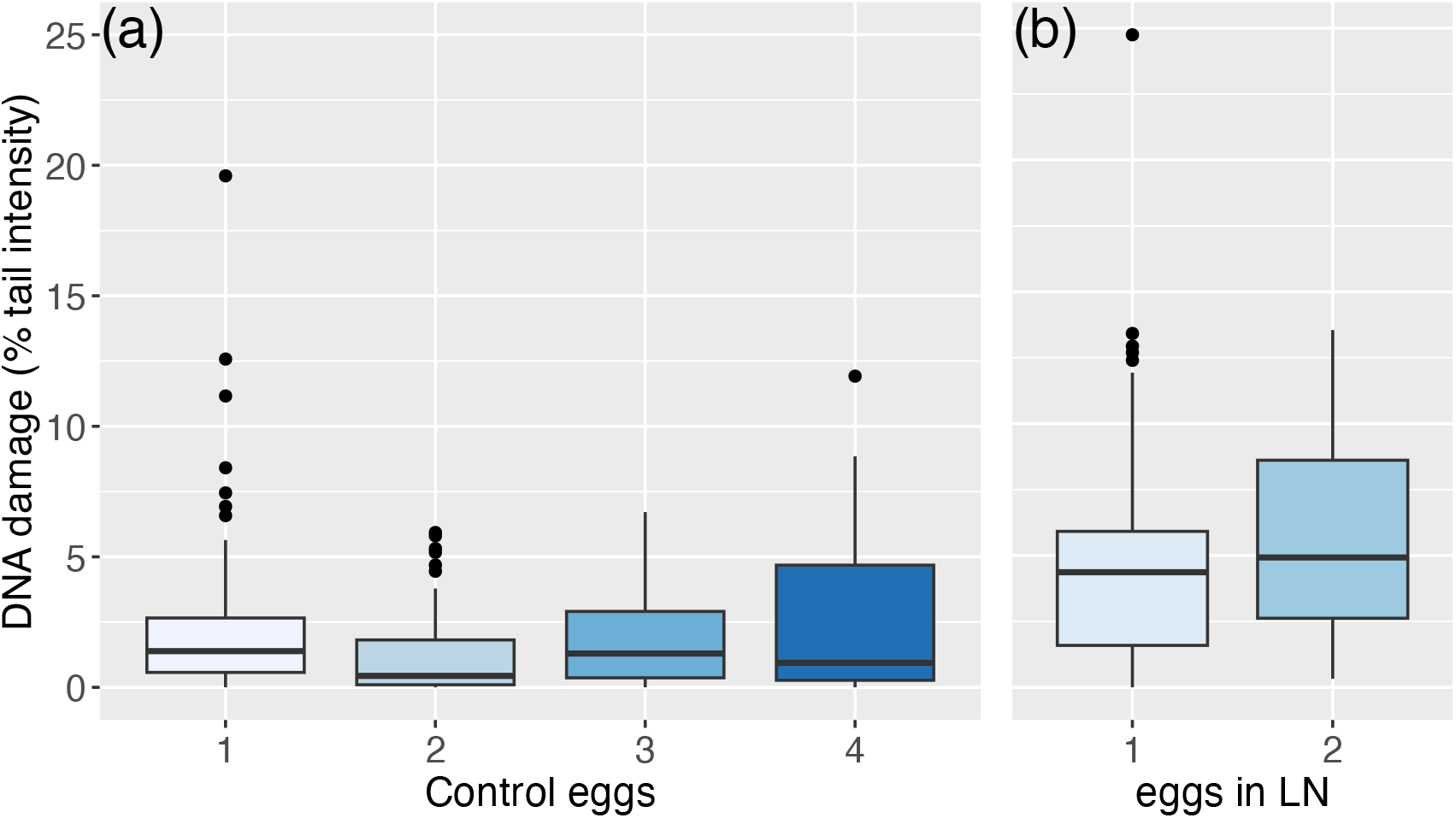
Single-egg CA tested on days-old dormant eggs of *Daphnia magna*: (a) four untreated eggs, (c) two eggs flash-frozen in LN. Boxplots show median, and 1st and 3rd quartile. Error bars represent minimum and maximum values, black circles represent outliers. DNA damage is measured as % tail intensity.

We additionally compared two *D. magna* eggs flash-frozen in LN and measured on separate slides (Fig. 2B). Here, median damage ranged from 4.37 to 4.93% and the Kruskal-Wallis test was not significant (Table S2, p=0.096).

### Freeze test

The impact of freezing was tested on single, days-old *Daphnia magna* eggs either frozen at -80°C or flash-frozen in LN (Fig. 3a). The damage in unfrozen control eggs was small (Table S1, median tail intensity 1.01%) while the DNA damage of eggs frozen in LN (median tail intensity 4.62%) and at -80°C (median tail intensity 13.01%) was significantly higher (Kruskal-Wallis rank sum test, p < 2.2e-16, Table S2). All comparisons were highly significant according to the Dunn test (Table S3, p < 0.0001).

**Fig. 3.**
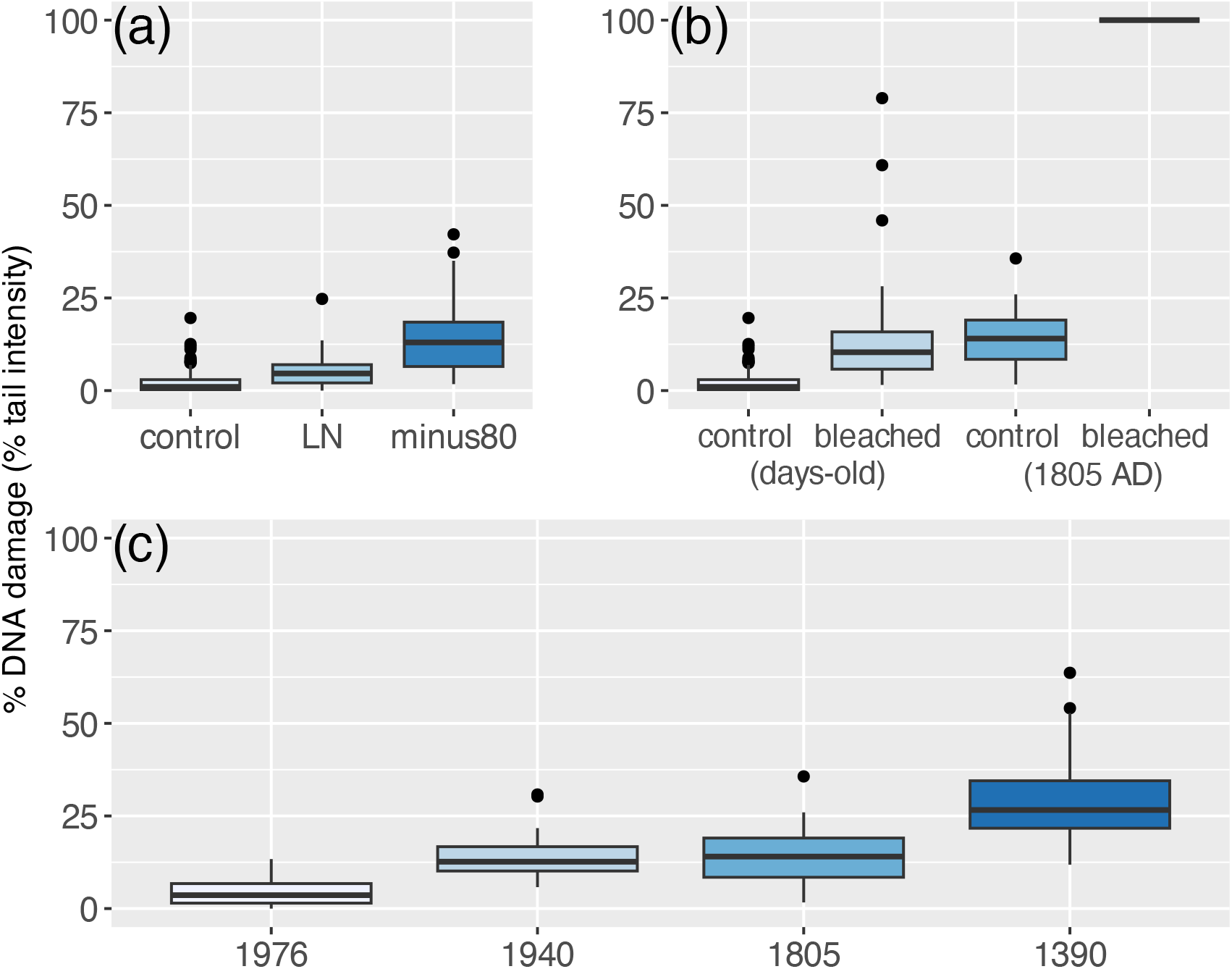
a-c. DNA damage (measured as % tail intensity) (a) on single dormant *D. magna* eggs (days-old) after 10-minute freezing (LN: Liquid Nitrogen, minus80: –80°C), control: no freezing), (b) after bleaching with a 5% industrial strength sodium hypochlorite solution on single dormant *D. magna* (days-old) and Arctic *D. pulicaria* eggs (historical, 1805 AD), (c) on single, historical dormant eggs of Arctic *D. pulicaria*, extracted from four sediment strata. Boxplots represent the median, and 1st and 3rd quartile. Error bars represent minimum and maximum values, black circles represent outliers.

### Bleach test

Days-old dormant eggs (*D. magna*) and historical, ca. 200-year-old dormant eggs (Arctic *D. pulicaria*) were treated with a 5% industrial strength sodium hypochlorite solution (Fig. 3b) and compared with unbleached controls. In the days-old eggs, DNA damage was significantly higher in bleached than unbleached eggs (Kruskal-Wallis rank sum test, p-value<2.2e-16, Table S2) with median values 1.01 and 10.36%, respectively (Table S1). In unbleached historical eggs DNA damage was 14%, but 100% in the historical eggs with no detectable nuclei present.

### Age test

The increase of DNA damage with increasing age of sedimentary *D. pulicaria* eggs was significant (Fig. 3c, Table S2, Kruskal-Wallis test, p < 2.2e-16) in eggs between 40 and 625 years old (median tail intensity 3.62% to 26.62%, respectively). All pairwise comparisons with the exception of that between eggs from 1940 AD and 1805 AD were highly significant (Table S3).

## Discussion

Our results show applicability of the CA protocol that we adapted for use with single *Daphnia* eggs (SE-CA), and additionally provide information on pre-treatment and storage of historical *Daphnia* eggs isolated from sediment archives as well as uncover the impact of egg age on DNA integrity.

While we show that the CA is applicable to individual eggs, careful pipetting is required to achieve satisfactory results. The main caveat is the extremely small size of the pellet obtained after homogenisation of individual eggs which becomes easily resuspended in the supernatant to be discarded. The limited number of cells in dormant eggs with an early gastrula stage embryo of ∼1000 cells (von Baldass 1941) results in a cell density in the initial LMA solution of about an order of magnitude lower (ca. 4 ×10^4^ cells/ml) than generally applied to conventional microscope slides (e.g. 2 × 10^5^ cells/ml, (Braafladt, Reipa & Atha, 2016). Further improvement of the method could be achieved by using minigels, such as a 12-gel slide which allow observation of as few as 200 cells per gel and thus could reduce the time spent on searching nuclei at low cell densities.

### Considerations for egg storage and for removal of exogenous DNA

Optimal storage of *Daphnia* eggs for molecular work, if further processing must be delayed, should focus on minimisation of further additional DNA damage. Application of the SE-CA after testing two freezing methods (LN and -80 C) revealed the highest DNA damage in eggs frozen at -80°C. DNA damage in LN-frozen eggs was only slightly higher than unfrozen eggs. The increase in cell damage after freezing human tissues at -80°C is thought to result from the formation of ice crystals due to slower freezing compared with LN cryopreservation at -196 °C (Bakhach, 2009). Freezing at -20°C results in decreased structural integrity in DNA molecules likely caused by a higher number of nicks in dsDNA (Chung *et al*., 2017), a process that could explain the higher damage also in the eggs tested here. Compatible with this, our results indicate that freezing in LN is the least destructive cryopreservation for *Daphnia* eggs.

Minimising exogenous DNA from sources of contaminant DNA on the surface of target organisms is important when preparing isolates for genomic applications. Bleaching with sodium chloride effectively damages DNA by oxidation and chlorination of nucleotides (Hayatsu, Pan & Ukita, 1971) and is commonly used for this type of decontamination prior to DNA analysis. For example, diluted bleach was used to decontaminate various invertebrate taxa prior to molecular gut-content analysis, including rotifers (Oh *et al*., 2020) and insects (Greenstone *et al*., 2012). Removal of exogenous DNA prior to Whole Genome Amplification (WGA) is especially important when working with historical, potentially damaged DNA because intact foreign DNA is likely to dominate in such samples. However, O’Grady et al. (2022) observed the failure of amplification of target DNA from historical *Daphnia* eggs after bleaching followed by WGA, suggesting that significant DNA damage is inflicted by this procedure. Here we showed that bleaching at relatively low concentrations already caused some DNA damage to days-old eggs but led to complete destruction of DNA in historical eggs. An undamaged egg membrane in the days-old eggs likely forms an effective barrier to bleach entering the egg. However, egg membranes damaged by aging can become permeable to bleach and enter the egg, destroying the contained DNA. Future microscopic analysis could shed light on egg membrane integrity to investigate the possibility of increased rupturing.

### DNA damage and egg age

Progressive DNA damage in ancient DNA results from both hydrolytic fragmentation by depurination, and deamination of cytosines, leading to sequencing artefacts (Orlando *et al*., 2021) or to inadequate DNA template for whole genome amplification that can result in overamplification of exogenous DNA (O’Grady *et al*., 2022). The DNA damage in days-old dormant eggs was ∼1% but > 20% in the ∼600-year-old egg. Inability to produce *Daphnia* DNA sequences via WGA of >300 year-old eggs from the same *D. pulicaria* egg bank (O’Grady *et al*., 2022) suggests that 20% DNA damage might already render the DNA inadequate for WGA procedures. For a more differentiated understanding of DNA damage beyond which genomic applications are likely to fail, a more detailed study is needed. The results obtained here suggest that the SE-CA can provide useful information on the likelihood of success for downstream genomic applications. Although destructive for tested material, it can give valuable information on general DNA preservation of other isolates from the same source (e.g. a single ephippium) or age range, and thus avoid unnecessary investment in time-consuming or costly procedures.

In conclusion, the CA is a cost-effective method that can be used in paleogenomics studies of egg or seedbanks, potentially of many taxa. Further studies could address a calibration of % tail intensity with actual quantity of DNA fragments (Azqueta & Collins, 2013) to gain better understanding of the thresholds of DNA damage for successful sequencing of whole genomes, or for resurrection of dormant eggs. Finally, we deliver a tool to quantify DNA damage in aquatic and terrestrial natural biobanks. Application of the SE-CA could be an important step toward a better understanding of the impact of pollution or of natural environmental factors such as salinisation or acidification on this invaluable historical resource.

## Acknowledgements

We thank John Coulbourne, University of Birmingham UK for providing the lab space for this work. DF received funding from the European Union’s Horizon 2020 research and innovation program under the Marie Skłodowska-Curie grant agreement No. 658714 and from the Deutsche Forschungsgemeinschaft (DFG, German Research Foundation) - 461099895. MW acknowledges Marie Skłodowska-Curie grant agreement No. 629892. We thank Caroline Sewell for providing *Daphnia magna* ephippia, and Maison Dane for help in the lab.

## Author contributions

Conceptualization: DF, MW; investigation: RS, AP, DF; writing - original draft preparation: RJ, DF; writing - review and editing: RS, AP, MW, DF.

