## Supplementary material for "Single-egg Comet Assay: a protocol to quantify DNA damage in aquatic dormant stages": 'Table S1' 'Table S2' 'Table S3'

**Table S1.** Statistical summary of results obtained for various assays testing the single-egg Comet Assay protocol developed here for dormant eggs of *Daphnia*. NA refers to no detectable nuclei present that could be scored.

| | **median** | ***min*** | ***max*** | ***nuclei scored (n)*** | ***% undamaged nuclei*** |
|---|---|---|---|---|---|
| *Single-egg test* | | | | | |
| slide (control) 1 | 1.39 | 0.00 | 19.60 | 60 | 8.33 |
| slide (control) 2 | 0.44 | 0.00 | 5.93 | 60 | 13.33 |
| slide (control) 3 | 1.30 | 0.00 | 6.71 | 63 | 12.70 |
| slide (control) 4 | 0.94 | 0.00 | 11.92 | 50 | 8.00 |
| slide (LN) 1 | 4.37 | 0.00 | 24.75 | 60 | 5.00 |
| slide (LN) 2 | 4.93 | 0.32 | 13.55 | 60 | 0.00 |
| *Freeze Assay* | | | | | |
| control | 1.01 | 0.00 | 19.60 | 233 | 10.73 |
| LN | 4.62 | 0.00 | 24.75 | 120 | 2.50 |
| -80°C | 13.01 | 1.75 | 42.18 | 60 | 0.00 |
| *Bleaching Assay* | | | | | |
| control_days-old | 1.01 | 0.00 | 19.60 | 233 | 10.73 |
| bleached_days-old | 10.36 | 1.53 | 78.93 | 51 | 0.00 |
| control_1805 AD | 14.04 | 1.66 | 35.67 | 50 | 0.00 |
| bleached_1805 AD | 100.00 | 100.00 | 100.00 | NA | 0.00 |
| *Age Assay* | | | | | |
| 1976 AD | 3.62 | 0 | 13.36 | 50 | 12 |
| 1940 AD | 12.65 | 5.785 | 30.79 | 50 | 0 |
| 1805 AD | 14.04 | 1.664 | 35.67 | 50 | 0 |
| 1390 AD | 26.62 | 11.86 | 63.64 | 50 | 0 |

**Table S2.** Kruskal-Wallis Rank Sum Tests testing the single-egg Comet Assay protocol developed here for dormant eggs of *Daphnia* and its application to various test related to storage, pre-treatment of eggs prior to molecular analyses, and the impact of egg age on DNA damage.

| | **Kruskal-Wallis $X^2$** | **df** | **p-value** | |
|---|---|---|---|---|
| Single-egg Test | | | | |
| *control* | 8.30 | 3 | 0.040 | * |
| *LN* | 2.77 | 1 | 0.096 | |
| Application tests | | | | |
| *Freeze Assay* | 161.95 | 2 | < 2.2e-16 | *** |
| *Bleaching Assay* | 173.15 | 2 | < 2.2e-16 | *** |
| *Age Assay* | 138.09 | 3 | < 2.2e-16 | *** |

**Table S3.** Pairwise comparison (Dunn test) of DNA damage (% tail intensity) in various assays testing the single-egg Comet Assay protocol developed here for dormant eggs of *Daphnia*. For details see text.

| | **Comparison** | **Z** | **p unadj** | **p adj** | |
|---|---|---|---|---|---|
| Single-egg Test | slide 1 : slide 2 | 2.656 | 0.008 | 0.047 | * |
| (days-old eggs) | slide 1 : slide 3 | 0.723 | 0.470 | 1.000 | |
| | slide 2 : slide 3 | -1.966 | 0.049 | 0.197 | |
| | slide 1 : slide 4 | 0.354 | 0.723 | 1.000 | |
| | slide 2 : slide 4 | -2.178 | 0.029 | 0.147 | |
| | slide 3 : slide 4 | -0.330 | 0.741 | 0.741 | |
| Freeze Assay | control : LN | -7.545 | 4.53E-14 | 6.79E-14 | *** |
| | control : -80°C | -11.876 | 1.58E-32 | 4.74E-32 | *** |
| | LN : -80°C | -5.512 | 3.55E-08 | 3.55E-08 | *** |
| Bleaching Assay | bleached_days-old - control_1805 | -0.950 | 3.42E-01 | 3.42E-01 | |
| | bleached_days-old - control_days-old | 9.508 | 1.94E-21 | 2.91E-21 | *** |
| | control_1805 - control_days-old | 10.645 | 1.85E-26 | 5.55E-26 | *** |
| Age Assay | 1309 AD : 1976 AD | 11.747 | 7.31E-32 | 4.39E-31 | *** |
| | 1309 AD : 1940 AD | 6.004 | 1.93E-09 | 3.85E-09 | *** |
| | 1976 AD : 1940 AD | 5.743 | 9.30E-09 | 1.39E-08 | *** |
| | 1309 AD : 1805 AD | 5.691 | 1.26E-08 | 1.51E-08 | *** |
| | 1976 AD : 1805 AD | 6.056 | 1.40E-09 | 4.19E-09 | *** |
| | 1940 AD : 1805 AD | 0.313 | 7.54E-01 | 7.54E-01 | |
